# Discrimination of Annonaceae using herbarium leaf reflectance spectra under limited sample size conditions

**DOI:** 10.1101/2025.09.02.673631

**Authors:** Khalil Boughalmi, Paola G. Santacruz Endara, Lily A. Bennett, Martin Ecarnot, Samantha Bazan, Denis Bastianelli, Laurent Bonnal, Thomas L.P. Couvreur

**Author notes:** Corresponding author : Khalil Boughalmi.

## Abstract

**Premise:** Herbarium collections offer an unparalleled archive of plant biodiversity, but their use for species identification through spectral data remains constrained by uncertain effects of preservation histories. This study assesses whether barium specimens can reliably predict species based on its leaf reflectance spectrum, despite variations in age, geographic origin, or conservation method under limited sample size conditions.

**Methods:** We scanned herbarium specimens of different ages and geographic distribution of 14 species of the pantropical Annonaceae. In addition, we used a second dataset of 9 species where some specimens were conserved in alcohol prior to drying and some not. We used five supervised classification models frequently used for high-dimensional data such as spectroscopy.

**Results:** All models achieved high accuracy (>80%) when trained on multiple specimens per species. However, when using only one specimen per species, accuracy varied substantially depending on the taxon.

**Discussion:** Our findings demonstrate that herbarium specimens often retain a strong taxonomic signal in their spectra, however, inter-individual variability affects accuracy in some taxa. These findings confirm the usefulness of herbarium spectroscopy as a non-destructive tool for species identification and offer a promising avenue for digitizing historical biodiversity data into high-dimensional trait space.

## INTRODUCTION

Herbaria hold hundreds of millions of preserved plant specimens, forming one of the largest and oldest repositories of biodiversity worldwide (Davis, 2023). These collections span more than four centuries of global plant exploration and constitute irreplaceable archives of taxonomic, geographic, and temporal information. Originally designed for taxonomy and floristics, herbaria have long been the foundation of plant classification and nomenclature. Yet, in recent decades, these collections are increasingly being reinterpreted as data-rich resources for ecology, evolution, and conservation biology (Besnard et al., 2018; Raxworthy and Smith, 2021). Herbarium specimens collected decades or even centuries ago are now used in trait-based ecology, functional biogeography, phylogenetics, and global change research (Nualart et al., 2017; Lang et al., 2019; Rosche et al., 2025), revealing new value in what was once considered passive documentation.

This shift has been catalyzed by the development of analytical tools capable of extracting meaningful data from historical plant material (Raxworthy and Smith, 2021; Davis, 2023). Among these, leaf reflectance, particularly near-infrared spectroscopy (NIRS), has emerged as a powerful and non-destructive technique to characterize plant functional traits (Cavender-Bares et al., 2017). Leaf reflectance spectra are a promising approach to species identification and even predicting functional traits (Kothari and Schweiger, 2022; Kothari et al., 2023) because they rapidly produce a snapshot of leaf chemical and structural composition. In such analyses, reflectance measurements typically span wavelengths from 350 to 2500 nm, with 400–700 nm corresponding to the visible range, 700–1100 nm to the near-infrared (NIR), and 1100–2500 nm to the shortwave infrared (SWIR). Reflectance is influenced by biochemical constituents such as cellulose, lignin, water, phenolics, and proteins, as well as by structural characteristics like leaf thickness or the presence of surface features such as hairs and stomata (Jacquemoud and Ustin, 2019).

Spectra therefore have the potential to serve as a rapid non-desctructive tool for plant species identification, as they may contain taxonomically informative features, sometimes referred to as the Species Spectral Signature (SSS) (Durgante et al., 2013). This taxonomic signal has led to a growing number of studies using reflectance spectra to discriminate between plant genera, species, and even populations. For example, Durgante et al. (2013) demonstrated that leaf reflectance spectra could successfully classify tropical tree species, while Meireles et al. (2020) revealed a phylogenetic component embedded in spectral variation at the angiosperm level. Other studies have confirmed the potential of NIRS for identifying hybrids, assessing intraspecific variability, and monitoring trait shifts across environments (Cavender-Bares et al., 2017; Stasinski et al., 2021).

Based on these observations, leaf reflectance spectroscopy has recently emerged as a promising tool for botanists, ecologists, and taxonomists alike. Furthermore, its non-invasive nature makes it particularly attractive for use on valuable or rare specimens, including historical herbarium material (types) including extinct species.

However, there is a critical caveat to this optimism. Most studies to date have relied on freshly collected, purpose-pressed leaves, prepared under relatively standardized and controlled conditions (referred to here as “pressed specimens”) (Durgante et al., 2013; Meireles et al., 2020; Kothari et al., 2023). Pressed specimens are not equivalent to herbarium specimens as they have not undergone mounting, long-term storage, transportation, or historical treatment. Herbarium specimens have often been exposed to a wide variety of in-field drying methods, conservation practices, and chemical treatments, many of which are undocumented, and all of which can potentially affect the chemical integrity of the leaves. The effects of these processes on leaf chemistry (and by extension, on spectral properties) are yet not well understood. This represents a fundamental limitation to the generalizability of existing findings.

For example, in humid tropical environments, alcohol is frequently used as a temporary preservative before heat drying, particularly during multi-day field expeditions (Hodge, 1947). While effective at preventing mold and tissue decay, alcohol is a strong solvent and oxidizer that can also impact DNA extraction (Forrest et al., 2019). Even today, practices such as applying glue, or drying methods vary considerably between herbaria and collectors. As a potential source of chemical modification, all these factors can introduce noise or bias into spectral data collected from herbarium sheets.

Moreover, the geographic origin of specimens introduces an additional layer of variation that reflects not only environmental conditions but also underlying genetic and phenotypic diversity within species. Climate, elevation, soil chemistry, and other environmental parameters influence leaf chemistry (Maire et al., 2015; Asner and Martin, 2016; Cao et al., 2020), while genetic variation, phenological stage, and ontogenetic differences can further contribute to spectral heterogeneity among specimens grouped under a single species definition. As a result, specimens of the same species, preserved differently and collected across distinct locations or developmental stages, may display substantial spectral divergence. If spectroscopy is to be used reliably for taxonomic or ecological inference on herbarium material, these multiple sources of variation must be understood and, where possible, accounted for within the analytical framework.

Despite these challenges, recent evidence suggests that herbarium specimens still retain valuable chemical and even strong taxonomic information (Neto-Bradley et al., 2025; White et al., 2025b). For instance, Foutami et al. (2018) have shown that the age of a specimen does not drastically affect some part of its chemical profile, even after decades of storage. While certain surface properties or volatile compounds may degrade over time, the core chemical constituents that drive spectral variation often remain stable (Mendes Resende et al., 2020). The key challenge is not whether herbarium spectra contain useful information, but whether this information is consistent, reproducible, and interpretable enough to enable species-level identification and classification, while remaining feasible at low sampling scales given the frequent scarcity of specimens per species and the logistical demands of projects involving hundreds or thousands of samples.

Here, we investigate several key questions regarding the application of spectroscopy to herbarium specimens. Specifically: (i) Can spectral data reliably support species-level identification? (ii) How many specimens per species are required to adequately represent intra-specific spectral variability? (iii) Are standard machine learning algorithms sufficiently robust to handle the heterogeneity typical of herbarium collections? and (iv) To what extent do pre-drying alcohol treatments affect the spectral signal used for classification? By addressing these questions, we aim to assess both the practical applicability and methodological limitations of herbarium spectroscopy under realistic working conditions.

To address the above questions, we focused on the pantropical plant family Annonaceae. With around 2500 species, it is one of the best-known tropical plant families in terms of taxonomy with global occurrence data and phylogenetic resources available (Couvreur et al., 2012, 2019; Nge et al., 2024). Despite this taxonomic maturity, species identification in Annonaceae can be challenging, particularly when reproductive structures (flowers or fruits) are absent or unavailable. This makes Annonaceae an ideal case study for evaluating the robustness of spectral approaches for species discrimination. Additionally, the family displays a high degree of ecological and biogeographic amplitude within the tropics, spanning environments from Amazonia to African rain forests and Southeast Asian regions, thus offering a natural gradient of variation in both environmental conditions and specimen collection practices (Pennington et al., 2004; Couvreur et al., 2011; Chatrou et al., 2012).

## METHODS

### Taxon sampling

We generated two datasets. First, to address the objective of species identification using herbarium-generated spectra, we scanned leaves from 14 Annonaceae species (**Table 1, Supplementary Table 1**) all stored in the herbarium of the Muséum national d’Histoire naturelle in Paris (P) (Le Bras et al., 2017). As is common for historical herbarium material, detailed information on pre-drying treatments or conservation practices was generally unavailable for these specimens, although such treatments are likely to have occurred. For each species we scanned a minimum of 9 specimens. The selected species were: 1) chosen from recently revised genera, ensuring expert-based identification; 2) represent different growth habits (liana, understory and canopy trees); 3) have a broad geographical distribution, including species from South America, Africa, Madagascar, and Southeast Asia; 4) have diverse geographical representation within species, including specimens from various countries and ecotypes when possible; and 5) are phylogenetically diverse across Annonaceae (at least one species for each of the four subfamilies, Nge et al. 2024). We also included two species pairs belonging to the same genera in *Ambavia* and *Hexalobus* (**Table 1, Supplementary Table 1**). In general, we prioritized flowering or fruiting specimens as these are better identified. This dataset is referred to as ID_PARIS.

**TABLE 1.** Overview of Paris herbarium specimens included in the study, with number scanned, geographic origin, specimen age span, and subfamily classification – ID_PARIS.

| Species | Specimens scanned | Geographic origin | Specimen age span | Subfamily |
| --- | --- | --- | --- | --- |
| *Ambavia capuronii*<br>(Cavaco & Keraudren)<br>Le Thomas | 17 | Madagascar | 1953-2013 | Ambavioideae |
| *Ambavia gerrardi* (Baill.)<br>Le Thomas | 18 | Madagascar | 1956-2010 | Ambavioideae |
| *Anaxagorea dolichocarpa* Sprague & Sandwith | 10 | French Guiana | 1845 - 2008 | Anaxagoreoideae |
| *Annona senegalensis*<br>Pers. | 9 | Zambia, Tanganyika, Mali, Sierra leone, Guinea | 1893 - 1951 | Annonoideae |
| *Anonidium mannii*<br>(Oliv.) Engl. & Diels | 11 | Cameroon | 1918 - 1995 | Annonoideae |
| *Duguetia calycina*<br>Benoist | 12 | French Guiana | 1857 - 2003 | Annonoideae |
| *Greenwayodendron suaveolens* (Engl. & Diels) Verdc. | 11 | Gabon, DR Congo, Central African Republic, Cameroon | 1881 - 1991 | Malmeoideae |
| *Hexalobus crispiflorus*<br>A.Rich. | 10 | Congo republic, Guinea, Cameroon, Gabon, Senegal | 1838 - 1988 | Annonoideae |
| *Hexalobus monopetalus*<br>(A.Rich.) Engl. & Diels | 11 | Mali, Cameroon, Central African Republic, Senegal, Niger, Ivory coast | 1929 - 2007 | Annonoideae |
| *Monanthotaxis enghiana*<br>(Diels) P.H.Hoekstra | 9 | Cameroon, Gabon, Central African Republic | 1948 - 1989 | Annonoideae |
| *Unonopsis guatterioides*<br>(A.DC.) R.E.Fr. | 12 | Brasil, Paraguay, Saint Vincent and the Grenadines, Surinam, Bolivia | 1886 - 1979 | Malmeoideae |
| *Uvaria grandiflora*<br>Roxb. ex Hornem. | 11 | Vietnam, Malaysia, China, Indonesia | 1823 - 2015 | Annonoideae |
| *Uvariastrum pierreanum*<br>Engl. | 10 | Gabon, Cameroon, Ivory coast, Congo republic, Liberia, Nigeria | 1948 - 2013 | Annonoideae |
| *Xylopia humblotiana*<br>Baill. | 9 | Madagascar | 1924 - 2007 | Annonoideae |

A second independent dataset was generated to examine the impact of alcohol preservation. For this we sampled nine species (**Table 2**) within the 50-ha Yasuní Forest Dynamics Plot (YFDP) located next to the Yasuní Research Station of the Pontificia Universidad Católica del Ecuador (Valencia et al., 2004). These species are the same as the study of Santacruz Endara et al. (2025) and cover the phylogenetic diversity of the family. First, specimens were sampled directly in the field at the Yasuní Research station of the Pontificia Universidad Católica del Ecuador (PUCE). Leaves were then pressed in newspaper (paper-flattened) and dried over an electric stove for 24 hours. For one or two specimens per species they were sprayed in 70-80% alcohol for three days, and then dried in an electric stove for 24 hours. This dataset is referred to as ID_YASUNÍ.

**TABLE 2.** Yasuní National Park dataset total specimen count and pre-drying alcohol-treated specimens – ID_YASUNÍ.

| Species | Total specimen count | Alcohol-treated specimen count |
| --- | --- | --- |
| *Anaxagorea brevipes* Benth. | 7 | 2 |
| *Duguetia hadrantha* (Diels) R.E.Fr. | 6 | 1 |
| *Guatteria guianensis* (Aubl.) R.E.Fr. | 6 | 1 |
| *Guatteria scalarinervia* D.R.Simpson | 5 | 1 |
| *Oxandra riedeliana* R.E.Fr. | 5 | 1 |
| *Tetrameranthus globuliferus* Westra | 7 | 2 |
| *Trigynaea triplinervis* D.M.Johnson & N.A.Murray | 6 | 1 |
| *Unonopsis veneficiorum* (Mart.) R.E.Fr. | 7 | 2 |
| *Xylopia cuspidata* Diels | 6 | 1 |

To place the two datasets into context, **Figure 1** summarizes the overall workflow, from specimen selection and scanning procedure acquisition to preprocessing, providing a framework for the methodological details that follow.

**Figure 1.**
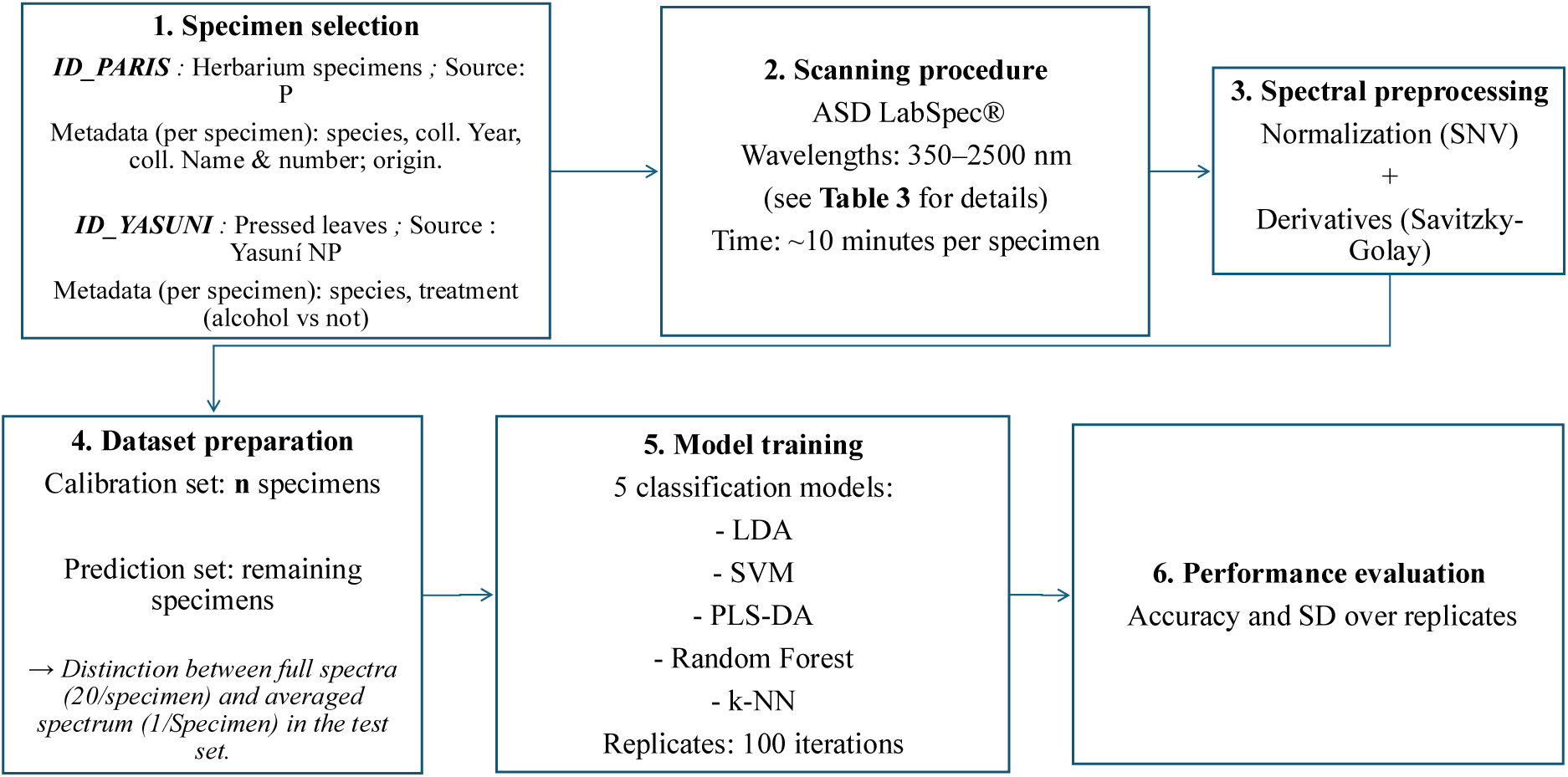
Workflow illustrating the classification pipeline used. Specimens from two datasets were scanned using spectrometers (ASD LabSpec), collecting 20 individual spectra per specimen. Spectral data was preprocessed, and databases were split into training and test sets across 100 replicates. Five supervised classification models were trained and then tested on either individual spectra or on spectra averaged per specimen. Model performance was assessed using accuracy.

### Spectral Measurements

Leaf specimens were scanned following a general protocol (Mersni et al., 2025), with dataset-specific adaptations (**Table 3**). In both datasets (ID_PARIS and ID_YASUNÍ), spectra were collected in the 350–2500 nm wavelength range using ASD LabSpec® instruments, with differences in probe configuration and acquisition setup between datasets. ID_PARIS measurements were acquired using a Rapid Analysis Probe coupled with a fiber optic illuminator, whereas ID_YASUNÍ spectra were collected using a 6 mm diameter pen probe. Spectra were generated by averaging multiple readings per scan (20 readings in ID_PARIS and 30 in ID_YASUNÍ), with 20 scans per specimen in ID_PARIS and 50 scans per specimen in ID_YASUNÍ. White reference and calibration procedures differed slightly between datasets (see **Table 3** for details). Measurements were performed by a single operator in ID_PARIS and by multiple operators in ID_YASUNÍ, but the same Spectral Black™ strip background was used.

**TABLE 3.** Comparison of spectral acquisition protocols and instrumentation between the ID_PARIS and ID_YASUNÍ datasets.

| Aspect | ID_PARIS | ID_YASUNÍ |
| --- | --- | --- |
| **Instrument** | ASD LabSpec® Standard-Res Lab Analyzer | ASD LabSpec® Pro spectrophotometer |
| **Probe type** | Rapid Analysis Probe (90° angle) + Fiber Optic Illuminator | 6 mm diameter pen probe |
| **Software version** | Indico Pro v6.5 | Indico Pro v3.1 |
| **Reference procedure** | White baseline references at start of session and every hour (Spectralon® white) | White reference every 15 minutes (Spectralon® Metal Spectral™) |
| **Black background** | Spectral Black™ strip beneath leaves | Spectral Black™ strip beneath leaves (mounted specimens) or larger black sheet (detached leaves) |
| **Sample count/scan** | 20 readings averaged to generate one spectrum | 30 readings averaged to generate one spectrum |
| **Integration time** | 68 ms<sup>-1</sup> | 68 ms<sup>-1</sup> |
| **Number of scans** | 20 scans per specimen | 50 scans per specimen |
| **Operator/time** | Single operator | Multiple operators |

In both datasets, spectra were collected on the adaxial side of mature leaves under controlled environmental conditions. Young or unfolding leaves were avoided, as well as the midrib region and areas affected by fungi, bryophytes, or glue. Whenever possible, multiple leaves per specimen were scanned, and measurements were distributed across the cleanest and least damaged areas. Care was taken to ensure contact between the probe and the leaf without applying excessive pressure that could damage the specimen.

### Data Processing

All data processing and analyses were conducted using R software (v4.4.1). The processing of spectral data ensures consistency, removes noise, and enhances relevant signals for further analysis and is composed of two steps. The first step involved applying a Standard Normal Variate (SNV) normalization (Barnes et al., 1989). For each spectrum, the mean and standard deviation of the 20 readings are calculated. Each reflectance intensity value is then normalized by subtracting the mean and dividing by the standard deviation, resulting in spectra with a mean of zero and unit variance. This standardization method removes a part of noise caused by external factors such as instrument settings, distance between the probe and the sample, or sample thickness. After normalization, the Savitzky-Golay filter (Savitzky and Golay, 1964) was used to compute the first derivative of the spectra. A polynomial order of 2 and a window size of 7 were used to smooth the data while capturing the rate of change at each wavelength. The first derivative accounts for variations in the slope of the spectra.

### Analytical Approaches

Five classification models were implemented to evaluate taxonomic discrimination: Partial Least Squares Discriminant Analysis (PLS-DA), Support Vector Machine (SVM), K-Nearest Neighbors (KNN), Linear Discriminant Analysis (LDA), and Random Forest (RF). Each model was selected for its already known capacity to handle spectral data sets (Kong et al., 2013; Trainor et al., 2017; Jombo et al., 2020; Wang et al., 2020). PLS-DA is widely used in chemometric and botanical applications due to its ability to model multicollinearity and extract latent components that maximize class separation (Richter et al., 2016; Song et al., 2018; Meireles et al., 2020; Kothari et al., 2023). LDA, while more limited in its assumptions, provides a simple and interpretable baseline model. Random Forest is a robust ensemble method capable of capturing non-linear relationships (Brabant et al., 2019). KNN offers a non-parametric approach that leverages local similarity in the spectral space and can be sensitive to subtle differences between specimens (Kong et al., 2013). Lastly, SVM has demonstrated strong performance in high-dimensional classification problems, particularly with spectral data, due to its capacity to define optimal decision boundaries in complex feature spaces, even when class separation is not linearly achievable (Brabant et al., 2019; Jombo et al., 2020). All models were trained and tested using stratified resampling with a strict separation by specimen identity to avoid data leakage, defined here as the presence of training or calibration specimens, or their associated spectra, in the test dataset. Specimen-level stratified sampling by species was applied in each iteration. For each species, six specimens were randomly assigned to a tuning subset. From the remaining specimens, approximately two-thirds were allocated to the calibration subset, and the remaining specimens formed the independent prediction subset. For PLS-DA, models were fitted using the plsda() function from the mixOmics package (Rohart et al., 2017). The number of components was optimized within each iteration by cross-validation on a dedicated tuning subset of the ID_PARIS dataset, testing values between 2 and 25. Hyperparameter optimization was conducted using five-fold cross-validation within the tuning subset (approximately 80% of specimens used for model fitting and 20% for internal validation at each fold). The optimal number of components was selected as the one minimizing cross-validated error rates. The SVM model was implemented using the caret interface to the e1071 (Meyer et al., 1999) and kernlab packages (Karatzoglou et al., 2024). Hyperparameter tuning was performed on the ID_PARIS dataset via cross-validation on a tuning subset, with a grid of sigma values (0.001, 0.01, 0.1) and cost values (0.1, 1, 10). Five-fold cross-validation was used within the tuning subset for hyperparameter selection. The best combination of sigma and C was retained for final calibration and testing. All predictor variables were standardized based on calibration data. For KNN, the knn3() function from the caret package (Kuhn et al., 2024) was used. The number of neighbors was tuned on the ID_PARIS dataset using cross-validation over k values from 1 to 10. Hyperparameter tuning was conducted using five-fold cross-validation within the tuning subset. Predictors were standardized according to calibration data, and the same transformation was applied to tuning and test sets. LDA was performed using the lda() function from the MASS package (Ripley et al., 2025). As LDA assumes uncorrelated predictors, a principal component analysis (PCA) was applied to the calibration data. The first set of components explaining at least 95% of the total variance were retained, and the same transformation was applied to tuning and test sets before fitting the LDA model. For Random Forest, the randomForest() function from the randomForest package (Breiman et al., 2024) was used with 500 trees (ntree = 500). All predictors with non-zero variance were included, and no additional feature selection or scaling was performed. Classification was based on majority vote across the ensemble of trees.

After hyperparameters were selected using the tuning subset, final models were trained exclusively on the independent calibration subset using the optimized parameters. The tuning subset was not reintegrated into the calibration phase, thereby maintaining strict separation between hyperparameter optimization, model fitting, and final evaluation. Model performance was then assessed on the independent prediction subset.

The models were applied to two distinct datasets: ID_PARIS, which was used for model training and testing with herbarium data, and ID_YASUNÍ, which was used to assess the effect of alcohol preservation on species discrimination. For ID_PARIS, the dataset was split into tuning, calibration, and prediction subsets, enabling full hyperparameter optimization. In contrast, analyses on ID_YASUNÍ and additional analyses on ID_PARIS at varying calibration sizes were conducted in a robustness-testing framework, where fixed hyperparameters were reused across all conditions. This ensured comparability across datasets and specimen sampling levels.

To simulate real-world conditions where the preservation method is generally unknown, alcohol-preserved specimens were randomly included in either the training or test dataset. This allowed us to evaluate how alcohol preservation might affect the models’ ability to distinguish between species. For each iteration, classification accuracy, or number of correct predictions divided by the total number of predictions, was calculated, and confusion matrices were pooled across resamples to evaluate model performance. To avoid bias, species were equally represented in all calibration datasets, irrespective of specimen availability. This balanced sampling ensured that species with more specimens did not dominate the models. Model predictions were compared to expert identifications to assess accuracy. Overall performance was computed for each model, and confusion matrices were used to highlight misclassification trends. We tested the models using both all available spectra per specimen (twenty spectra) and their average. To further explore the limitations of using a single herbarium specimen to represent a species, we conducted an analysis using the ID_PARIS dataset. We first evaluated model performance with only one randomly selected specimen per species, using its twenty spectra for training and the spectra of all remaining specimens for testing. We then extended the analysis by progressively increasing the number of training specimens from one to eight, using all their spectra for training and the spectra of the remaining specimens for testing.

Finally, to ensure robustness and consistency, all analyses were validated using 100 randomly generated iterations. This repeated validation allowed us to assess the stability of model performance across different sampling conditions and to ensure reliable results.

## RESULTS

### Specimen spectrum sampling

A total of fourteen species were scanned in the P herbarium. Using the previously described workflow (**Fig. 1**), we were able to scan up to 18 specimens for some species. Preparing the specimen, entering the required parameters, and initiating the first scan takes approximately 8 minutes. Once this setup is completed, acquiring the remaining scans is rapid, so that collecting a total of 20 spectra requires about 10 minutes per specimen. This enabled us to scan 160 specimens over 5 days (see **Table 1 & Table S1** for details). The scanned specimens ranged in age from 9 to 201 years, with an overall average of 69 years. Some species include older specimens (e.g., *Annona senegalensis*: mean 107 years, SD = 23; *Uvaria grandiflora*: mean 89 years, SD = 53; *Anaxagorea dolichocarpa*: mean 88 years, SD = 64), while others include younger ones (e.g., *Xylopia humblotiana*: mean 30 years, SD = 22; *Ambavia spp.*: mean 33 years, SD = 15) (**Fig. S1**). Several issues arose during sampling due to specific conditions encountered with some specimens, such as glue or highly curled leaves. Among the 160 specimens, less than five were affected by the scanning process, and the impacts were never severe and generally consist of leaf tears on curled specimens and leaf detachment. In the case of the samples from Yasuní, the scanning workflow was much more efficient because the data had already been collected and the leaves were not fixed in an herbarium voucher. Despite taking more scans per specimen (50 instead of 20), it took less than five minutes to measure a whole specimen.

### Spectrum discrimination using herbarium specimen

For all five classification models tested, we observed an overall strong capacity to discriminate herbarium specimens at the species level. Using all spectra from training specimens species and all spectra (measurement-level) from the remaining specimens for testing, we obtained the following overall accuracies: 84.74% (SD = 5.23%) for LDA, 81.45% (SD = 4.50%) for SVM, 81.06% (SD = 4.47%) for RF, 78.57% (SD = 4.77%) for PLS-DA, and 75.63% (SD = 4.08%) for KNN (**Fig. 2A**). When each specimen in the test set was represented by a single averaged spectrum rather than the full set of 20 individual spectra (specimen-level), the discrimination capacity were even higher for all models: 87.48% (SD = 5.57%) for LDA, 86.35% (SD = 4.47%) for SVM, 84.96% (SD = 5.60%) for RF, 83.03% (SD = 5.82%) for PLS-DA, and 79.23% (SD = 4.86%) for KNN( **Fig. 2A**). This remained true independent of species diversity in terms of geographic origin or collection date (**Fig. 2B**). However, two species from the same genus, *Hexalobus crispiflorus* and *H. monopetalus*, appeared to be less accurately classified, with misclassifications from one to the other. In most models using individual spectra as the test, there was a tendency for spectra to be classified as the other species, with a slight bias toward misclassification in favor of *H. crispiflorus*. When using the average spectrum per specimen in the test set, this pattern persisted, where *H. monopetalus* was more frequently predicted as *H. crispiflorus* than the reverse (**Fig. 2B**). This trend was especially pronounced in LDA, where nearly all *H. crispiflorus* specimens were correctly classified (79.50%) but only 66.67% of *H. monopetalus,* the misclassified ones were all predicted as *H. crispiflorus*. The two *Ambavia* species showed a better discrimination rate without a distinct four-square misclassification pattern. However, there was a slight but consistent tendency to misclassify *A.capuronii* as *A. gerrardii*. *Uvaria grandiflora* also appeared to be less well classified, where its correct classification rate was among the lowest observed. Overall, despite a general increase in model accuracy when using the average spectrum, the same species tended to be misclassified across both approaches and generally in the same predicted species (**Fig. 2B**).

**Figure 2.**
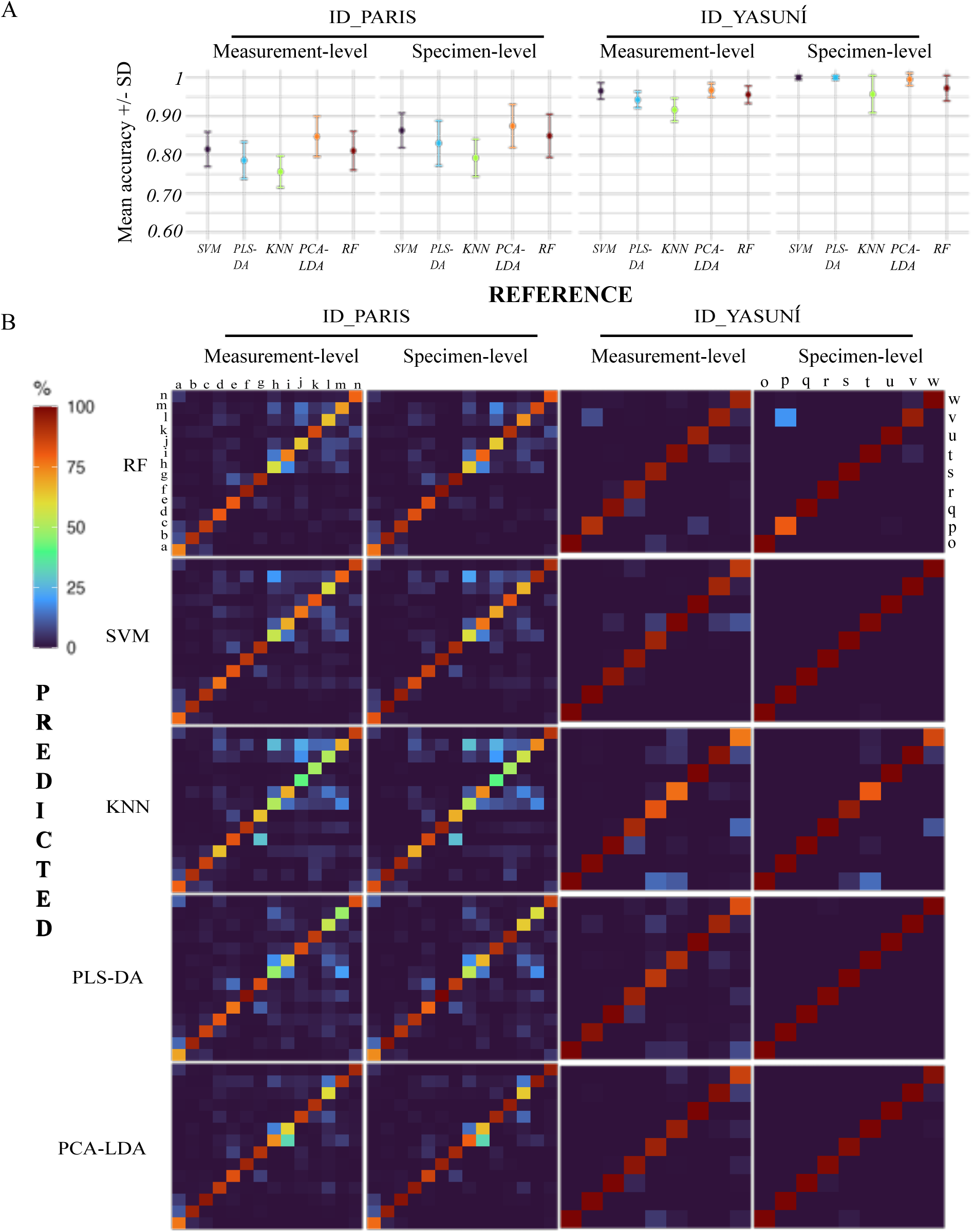
Species-level classification accuracy of leaf spectra. Classification performance for 14 species from ID_PARIS (a-*Ambavia capuronii*, b-*Ambavia gerrardi*, c-*Anaxagorea dolichocarpa*, d-*Annona senegalensis*, e-*Anonidium mannii*, f-*Duguetia calycina*, g-*Greenwayodendron suaveolens*, h-*Hexalobus crispiflorus*, i-*Hexalobus monopetalus*, j-*Monanthotaxis enghiana*, k-*Unonopsis guatterioides*, l-*Uvaria grandiflora*, m-*Uvariastrum pierranum*, n-*Xylopia humblotiana*) and 9 species from ID_YASUNÍ (o-*Anaxagorea brevipes*, p-*Duguetia hadrantha*, q-*Guatteria guianensis*, r-*Guatteria scalarinervia*, s-*Oxandra riedeliana*, t-*Tetrameranthus globuliferus*, u-*Trigynaea triplinervis*, v-*Unonopsis veneficiorum*, w-*Xylopia cuspidata*) using five models: Partial Least Squares Discriminant Analysis (PLS-DA), k-Nearest Neighbors (KNN), Principal Component Analysis followed by Linear Discriminant Analysis (PCA-LDA), Support Vector Machines (SVM), and Random Forest (RF) from 100 independent replicates using 75 % randomly selected specimens in training (6 for ID_PARIS, 3 for ID_YASUNÍ) all the others in the test, Measurement-level column mean that all specimen spectra where classify and Specimen-level column that only the average spectrum is classified. **(A)** Overall model accuracy and standard deviation. **(B)** Confusion matrices represent mean classification results. Values indicate the proportion of spectra predicted for each species class.

### Alcohol treatment effect

All models showed an accuracy above 90% when every spectrum was included in the test set, and no model accuracy was below 95% when the species average spectrum was used in the test set. Three models (SVM, PLS-DA and LDA) reached classification rates above 99%, with very low standard deviation (SVM and PLS-DA <0.005; LDA <0.02 (**Fig. 2A**)), indicating nearly all specimens were correctly classified to their species, regardless of whether alcohol-treated specimens were included in the training or test datasets. We did not observe a strong genus-level misclassification pattern, similar to what was seen in *Hexalobus*, among the two *Guatteria* species included in this dataset especially with the average specimen in the test dataset (**Fig. 2B**). Additionally, we found that alcohol-preserved specimens do not appear to be distant in the 3D space defined by the first three PCA components of the average spectrum for each specimen (**Fig. S2A**). These specimens cluster closely with others from the same species and are never misclassified into the spectral space of a different species. The average raw spectra themselves also generally share the same overall shape as those of other non-alcohol-preserved specimens from the same species, while differences in magnitude can be observed (**Fig. S2B**).

### One specimen to represent them all

We observed substantial variability in classification accuracy depending on the species when training the models with only a single specimen but using its 20 spectra (**Fig. 3**). For some species, such as *Anaxagorea dolichocarpa*, *Duguettia calycina* or *Ambavia gerrardii*, accuracy remained high more than 80% across all models (Mean values *A. dolichocarpa* 85.22; *A. gerrardii* 82.77; *D. calycina* 84.64 (**Fig. 3**)), while for others like *Uvaria grandiflora* or both *Hexalobus* species, it dropped below 50% (Mean values *H. monopetalus* 49.33; *H. crispiflorus* 43.47; *U. grandiflora* 37.64 (**Fig. 3**)). Despite this variability, model performance tended to follow similar patterns: species that were generally well classified remained so across models, and the same held for poorly classified species. Nonetheless, LDA generally outperformed the other methods, showing the highest classification accuracy in 12 out of 14 species. In certain cases, this performance gap was particularly marked (e.g. *Monanthotaxis enghiana*).

**Figure 3.**
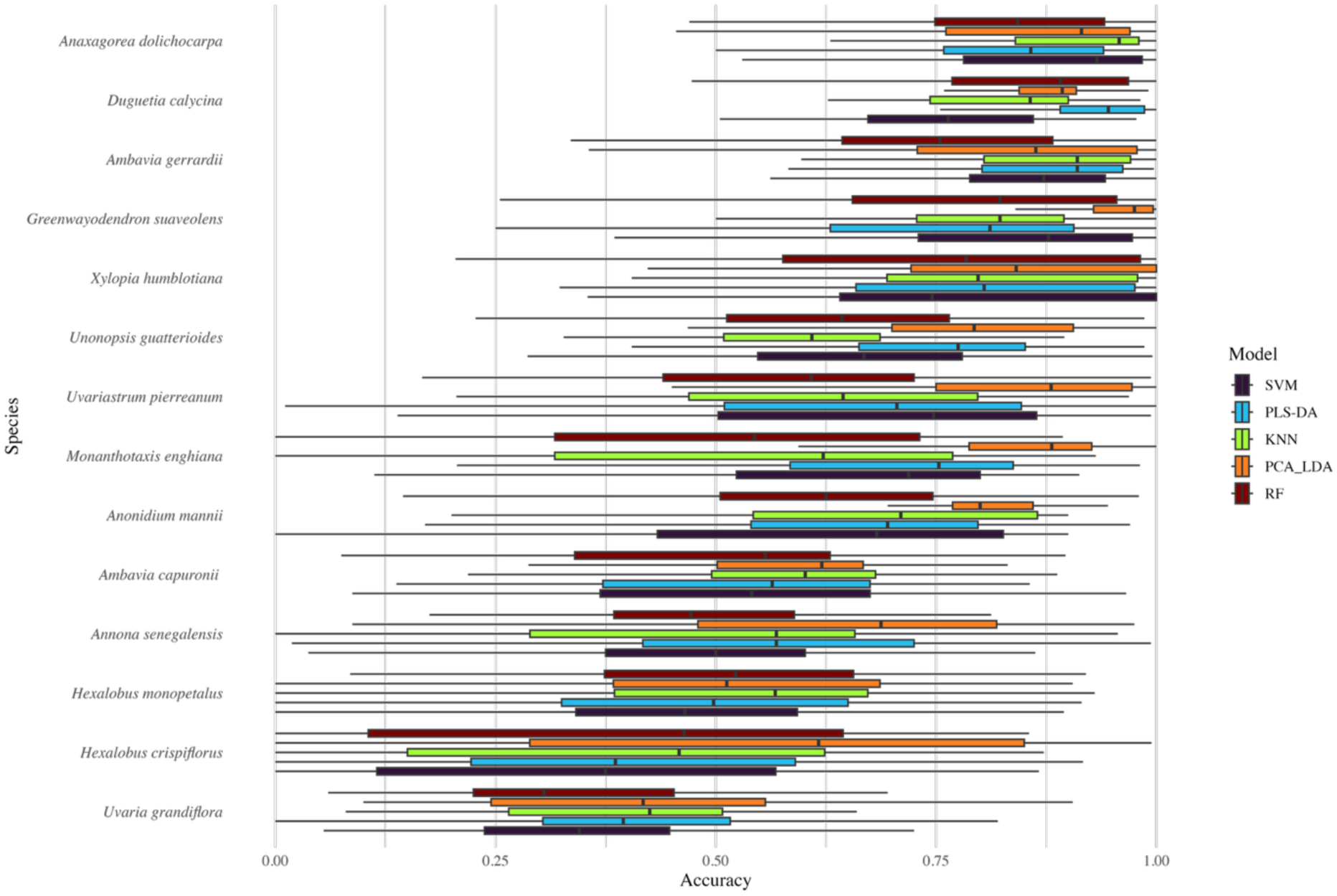
Performance of models trained on a single specimen per species. Classification accuracy (boxplots) across 100 replicates for each species and model, using one randomly selected specimen (20 spectra) as the training dataset. Background tiles indicate the mean classification accuracy across all models for each species. Results are based on the ID_PARIS dataset. Species are ordered from highest (top) to lowest (bottom) mean accuracy.

With an average accuracy of 65.4% (**Fig. 4A**), the models showed an important improvement, reaching 78.2% with only two specimens used for training, and stabilizing near 90% from five specimens onward, with only slight increases beyond that point (**Fig. 4B**). LDA consistently outperformed the other models at all specimen counts, achieving over 70% accuracy with just one specimen and reaching more than 90% from three specimens onward.

**Figure 4.**
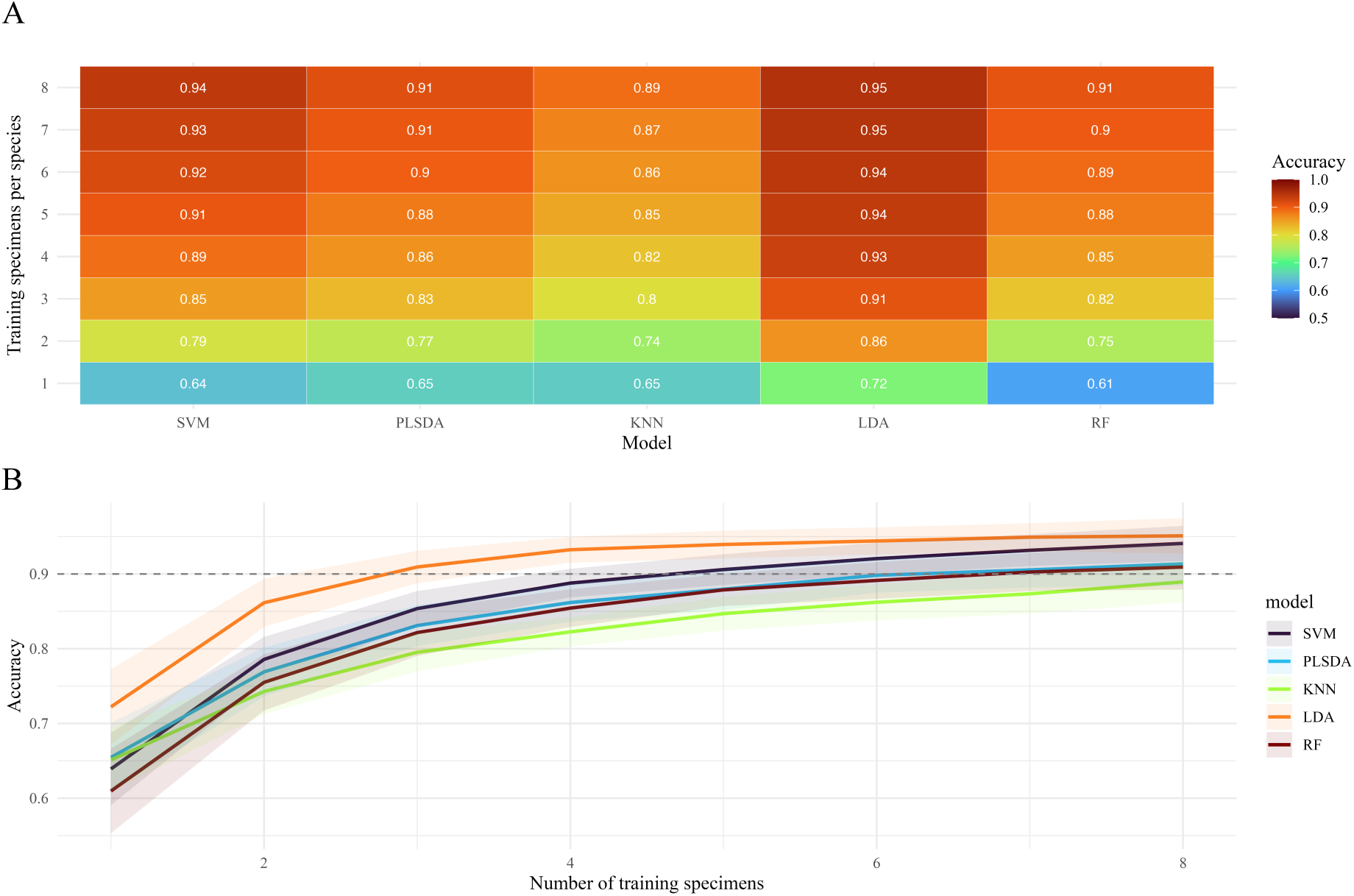
Effect of the number of training specimens per species on classification accuracy. **(A)** Heatmap of mean classification accuracy across 100 resampling iterations for each model and training size (1 to 8 specimens per species) using the ID_PARIS dataset. **(B)** Accuracy trajectories (± standard deviation) across the same iterations, showing model performance as a function of the number of training specimens per species.

## DISCUSSION

We demonstrate that herbarium leaf spectra contain sufficient information to enable accurate species classification, even with small sample sizes and samples that are decades old. Across most taxa tested, classification performance remained consistently high (**Fig. 2A**), supporting the idea that herbarium material can preserve relevant traits for classification (Cavender-Bares et al., 2017; Meireles et al., 2020; Kothari et al., 2023; Neto-Bradley et al., 2025; White et al., 2025b). Importantly, high accuracy (often >90%) was achieved using as few as five specimens per species for model training (**Fig. 4**). Even under single-specimen conditions, performance far exceeded random expectations (**Fig. 3**), indicating that the spectral signature (Durgante et al., 2013) of many species is both coherent and stable.

Classification success was not universal across taxa, and performance varied among species. In the two instances of congeneric species included in this study, classification accuracy was reduced compared to intergeneric discrimination, yet remained well above random expectations. This was particularly evident for the two *Hexalobus* species, which showed comparatively lower accuracy across models (**Fig. 2B**) but were still discriminated at rates substantially higher than chance. These results support the idea that closely related species present greater challenges for spectral discrimination, even when they are morphologically, ecologically, and phylogenetically distinct (Botermans et al., 2011; Dagallier et al., 2023). For example, *Hexalobus crispiflorus* and *H. monopetalus* occupy contrasting environments, rainforest and savanna, respectively, yet their spectral signatures did not reflect this ecological divergence. Such cases indicate that while reflectance spectroscopy is generally effective, it may occasionally fail to capture biologically meaningful differences among certain closely related species. In contrast, *Greenwayodendron suaveolens*, a widespread generalist tree species in the Congo Basin (Lissambou et al., 2018), was robustly classified despite spanning multiple countries and collection periods (**Fig. 3**), highlighting the potential of spectroscopy to detect stable traits even under broad ecological and geographic variation. It is noteworthy that misclassification was not restricted to congeneric species. *Uvaria grandiflora*, which was the sole representative of its genus in the dataset, also showed reduced classification performance, indicating that factors beyond taxonomic proximity, such as intraspecific variability, specimen condition, or unobserved preservation artefacts, may contribute to classification difficulty. An important consideration is that most species included in this study belong to different genera, validating this approach at the higher taxonomic level. However, species-level discrimination in more closely related species may require larger training sets.

We evaluated five classification models (PLS-DA, LDA, RF, KNN, and SVM), each selected for its capacity to handle high-dimensional spectral data (Kong et al., 2013; Trainor et al., 2017; Jombo et al., 2020; Wang et al., 2020). Although all models achieved strong performance, they differed in robustness and scalability (**Fig. 2–4**). LDA emerged as the most accurate method in this study, likely benefiting from the structured nature of the dataset and prior dimensionality reduction. It is nevertheless worth noting that, despite its strong overall performance, LDA also exhibited limitations when discriminating among congeneric species, highlighting the increased difficulty of intra-generic classification. SVM also performed consistently well, reflecting its strength in handling complex decision boundaries in high-dimensional space. In contrast, RF and KNN showed greater variability, while PLS-DA, although effective, is mathematically less suited to classification as the number of classes increases (Trainor et al., 2017). Accuracy improved with increasing training set size but plateaued beyond five specimens per species (**Fig. 4**), a promising result for specimen-poor but species-rich groups.

We deliberately retained the full spectral range for model training, rather than automatically excluding the first and last wavelength regions as commonly recommended (e.g. Meireles et al., 2020), in order to explore the complete spectral signal present in herbarium specimens. We also used spectra that, as shown in **Fig. S2** for ID_YASUNÍ and **Fig. S3** for ID_PARIS, occasionally display abrupt jumps in reflectance related to instrument characteristics. In our case, these artefacts did not appear to reduce model performance when using processed spectra. Consequently, the classification performance reported here may represent a conservative estimate and could potentially be further improved through targeted wavelength selection or trimming, even though accuracy was already high.

In addition to overall classification performance, this study represents a first step toward exploring the effect of alcohol preservation prior to drying on herbarium-based spectral classification. Alcohol preservation is commonly used during specimen preparation, particularly in humid tropical environments, where it helps prevent tissue degradation, mold development, and other forms of contamination. Despite its widespread use, quantitative information on the proportion of alcohol-preserved specimens in herbarium collections, as well as on exposure duration and concentration, remains unavailable. It is nevertheless reasonable to assume that a substantial fraction of tropical herbarium specimens has been exposed to alcohol at some stage of preparation, in some cases for extended periods.

Alcohol preservation may influence dried herbarium specimens through both physical and chemical mechanisms. Physically, alcohol can induce tissue shrinkage or structural alteration, while chemically it is a well-known solvent for chlorophyll and other compounds contributing to leaf optical properties. In our experimental setup, visual inspection of PCA space based on averaged spectra indicates that alcohol-treated specimens generally occupy species-specific regions overlapping with those of untreated specimens (**Fig. S2A**). Although slight shifts in spectral magnitude were observed (**Fig. S2B**), the overall spectral shape and the main taxonomic signal appeared to be preserved.

Consistent with these observations, species-level classification accuracy remained high when alcohol-treated specimens were included in either the training or test datasets. Across all models, accuracy exceeded 95%, with particularly strong performance for SVM, LDA, and PLS-DA (**Fig. 2A**), indicating that alcohol-treated specimens retained sufficient spectral information for accurate species discrimination in this experimental context. However, several aspects of the design should be considered when interpreting these results. The alcohol dataset comprised a relatively limited number of specimens and species, which likely facilitated discrimination. In addition, classification success may partly reflect the behavior of supervised learning algorithms, which can downweight spectral regions most affected by chemical alteration and rely on more stable wavelength ranges for discrimination. Taken together, these results suggest that under controlled, short-term exposure conditions and at a limited taxonomic scale, alcohol-preserved herbarium specimens can retain a usable taxonomic signal for spectral classification. Rather than providing a generalized assessment of alcohol preservation effects, this experiment offers an initial indication that moderate chemical alteration does not necessarily preclude the use of herbarium spectra in supervised classification frameworks.

More broadly, pre-drying preservation methods, including those involving alcohol, vary widely across herbarium collections and are often insufficiently documented. Differences in exposure duration (i.e. several weeks), concentration, and storage conditions are likely to produce a wide range of chemical alterations, some exceeding those captured by controlled experiments. Addressing this variability will require dedicated reference datasets, either based on specimens with well-documented preparation histories or purpose-designed experiments explicitly testing preservation treatments. While recent studies have focused on specimen aging and drying methods (Kühn et al., 2024), chemical treatments, aside from glue-related effects (White et al., 2025b), remain comparatively understudied, despite their potential influence on spectral integrity.

While these results are encouraging, our study was conducted under controlled conditions using specimens from a single herbarium. We did not isolate the effects of specimen age or geographic origin, reflecting the reality that such metadata are often incomplete or unavailable. By working under realistic constraints, our goal was to assess model performance in conditions commonly encountered in herbarium-based research, balancing methodological rigor with operational feasibility.

From a methodological perspective, these challenges raise important questions regarding specimen usability, potential exclusion thresholds, and analytical strategies that explicitly account for preservation-related effects. Future developments may include preservation-aware preprocessing pipelines, model architectures accommodating chemically induced spectral shifts, or scan-level approaches that retain information currently lost through spectrum averaging. A related methodological consideration concerns the decision to use a single measurement per specimen versus averaging multiple scans. While averaging replicate scans can reduce instrumental noise and improve classification stability, it may also obscure within-specimen variability that could be biologically meaningful. The optimal strategy likely depends on the research objective, whether prioritizing classification accuracy, trait estimation, or the study of intra-individual variation. Artefacts introduced by glue, fungal contamination, or leaf structure further complicate interpretation. Although preprocessing methods such as SNV and Savitzky–Golay filtering help mitigate noise, they do not fully resolve the boundary between noise and true biological variation. The choice of preprocessing methods should therefore be preceded by systematic testing to determine which treatments are most effective and discriminative, ideally by evaluating multiple combinations of preprocessing steps.

Beyond analytical considerations, the practical implementation of spectral acquisition also deserves attention. Using our methods, scanning required approximately 10 minutes per specimen, allowing up to 50 specimens to be processed during a standard 8-hour working day. This throughput highlights the operational feasibility of herbarium-based spectral acquisition. However, more detailed protocols are now available (White et al., 2025a), and the time required may vary depending on the objectives of the research question, the level of metadata recorded, and the degree of replication implemented.

These practical considerations further emphasize the importance of harmonized protocols when combining datasets across studies or collections. Differences in specimen composition, preservation history, sensor characteristics, and measurement conditions may introduce systematic biases if datasets are pooled uncritically. In particular, instrument-specific artefacts, such as abrupt reflectance discontinuities or jumps between sensor regions **(Fig. S2-S3**), can vary among devices and may affect spectral continuity in ways that are not immediately evident. Such features should be carefully evaluated and, where necessary, corrected or accounted for before merging datasets acquired with different instruments. Careful dataset-specific evaluation and harmonization are therefore essential prerequisites for meaningful data aggregation.

More broadly, harmonizing protocols across studies remains essential for building interoperable spectral datasets. Initiatives such as IHerbSpec (Cavender-Bares et al., 2025) provide a strong foundation by promoting standardized specimen preparation, scanning environments, and metadata reporting (White et al., 2025a). Adoption of such frameworks will be key to ensuring that herbarium spectral data are both locally robust and globally reusable.

## Conclusion

The application of near infrared spectroscopy to herbarium specimens is still in its infancy. Our results show that even old specimens retain enough taxonomic signal in their leaf reflectance spectra to allow accurate species classification using standard machine learning approaches. This confirms spectral scanning as a valuable, non-destructive source of data for studies in taxonomy, ecology, and evolution (Meireles et al., 2020; Stasinski et al., 2021; Kothari and Schweiger, 2022).

The potential of spectroscopy applied to herbarium-preserved specimens opens exciting perspectives for plant biodiversity research, adding yet another important dimension to the use of herbaria (Davis 2023). In addition to complementing traditional taxonomic research, spectroscopy could also inform leaf functional traits and, in combination with advances in machine learning (Soltis et al., 2020), enable large-scale, non-destructive trait estimation, species identification, and even the detection of cryptic diversity across historical collections. Ensuring data quality and expanding spectral libraries across taxa and regions will be key to unlocking the full potential of this approach.

## CRediT Author contributions

**KB**: Conceptualization, Data curation, Formal analysis, Writing (original draft). **TLPC**: Conceptualization, Supervision, Methodology, Validation, Writing (review & editing), Investigation (field sampling), Funding acquisition. **PGSA**: Methodology, Investigation (field sampling), Writing (review) **LAB**: Methodology, Writing (review & editing). **ME**: Methodology, Writing (review & editing). **SB**: Methodology (instrumentation), Writing (review). **DB**: Methodology (instrumentation), Writing (review). **LB**: Methodology (instrumentation), Writing (review).

## Acknowledgment

The authors thank the Muséum national d’Histoire naturelle (MNHN), Paris, for access to its herbarium collections, and the Yasuní National Park authorities for permitting field sampling and supporting research. We are grateful to Dr. Laura Holzmeyer for taxonomy training prior to specimen selection at P. This research was supported by the European Research Council (ERC) under the European Union’s Horizon 2020 program (grant agreement No. 865787).

## Data - Code availability

All code and data used in this study are available from the IRD Forge repository : https://forge.ird.fr/diade/global_erc/boughalmi_nirs_paris_yasuni_data_2025

## SUPPLEMENTARY

**Table S1.**
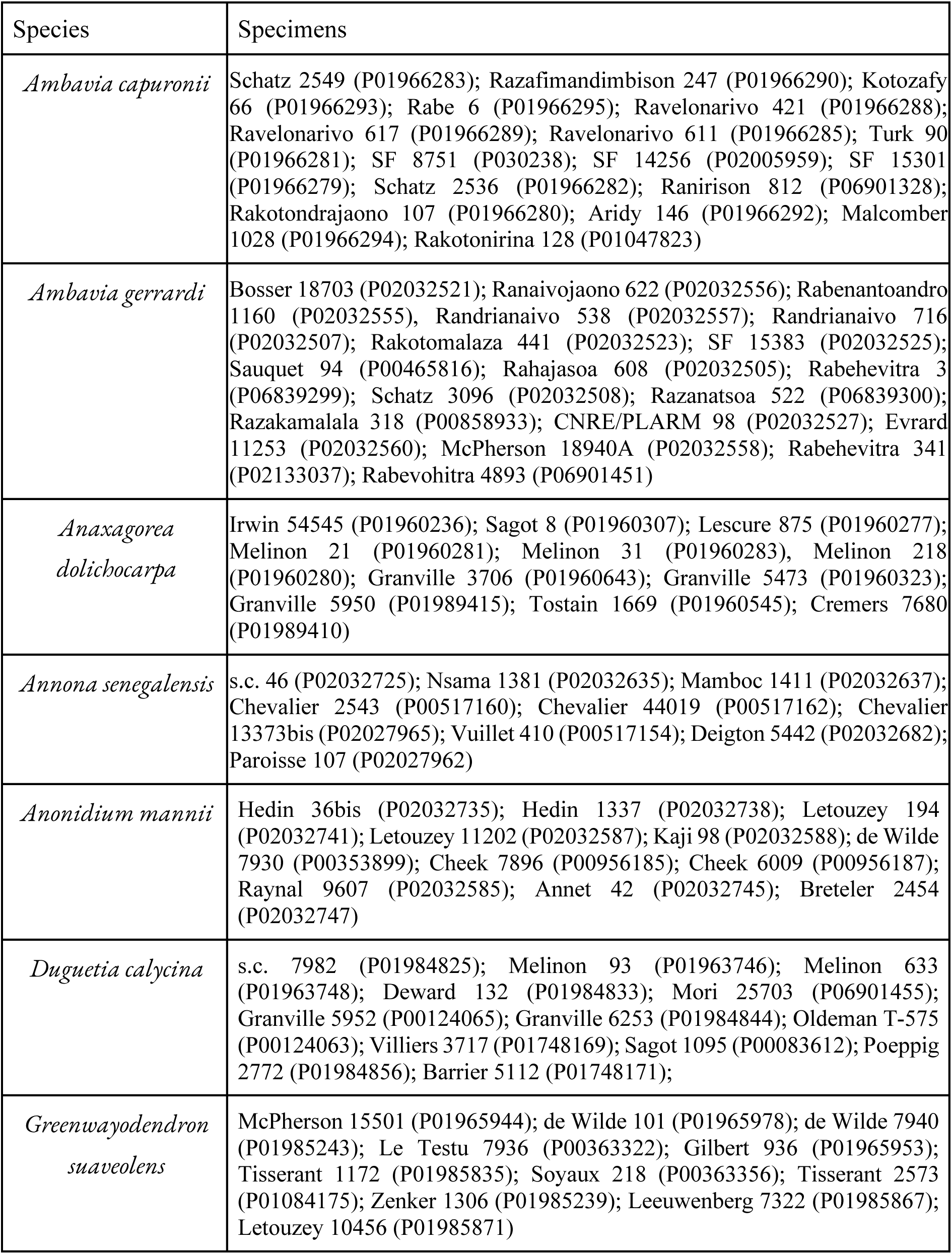
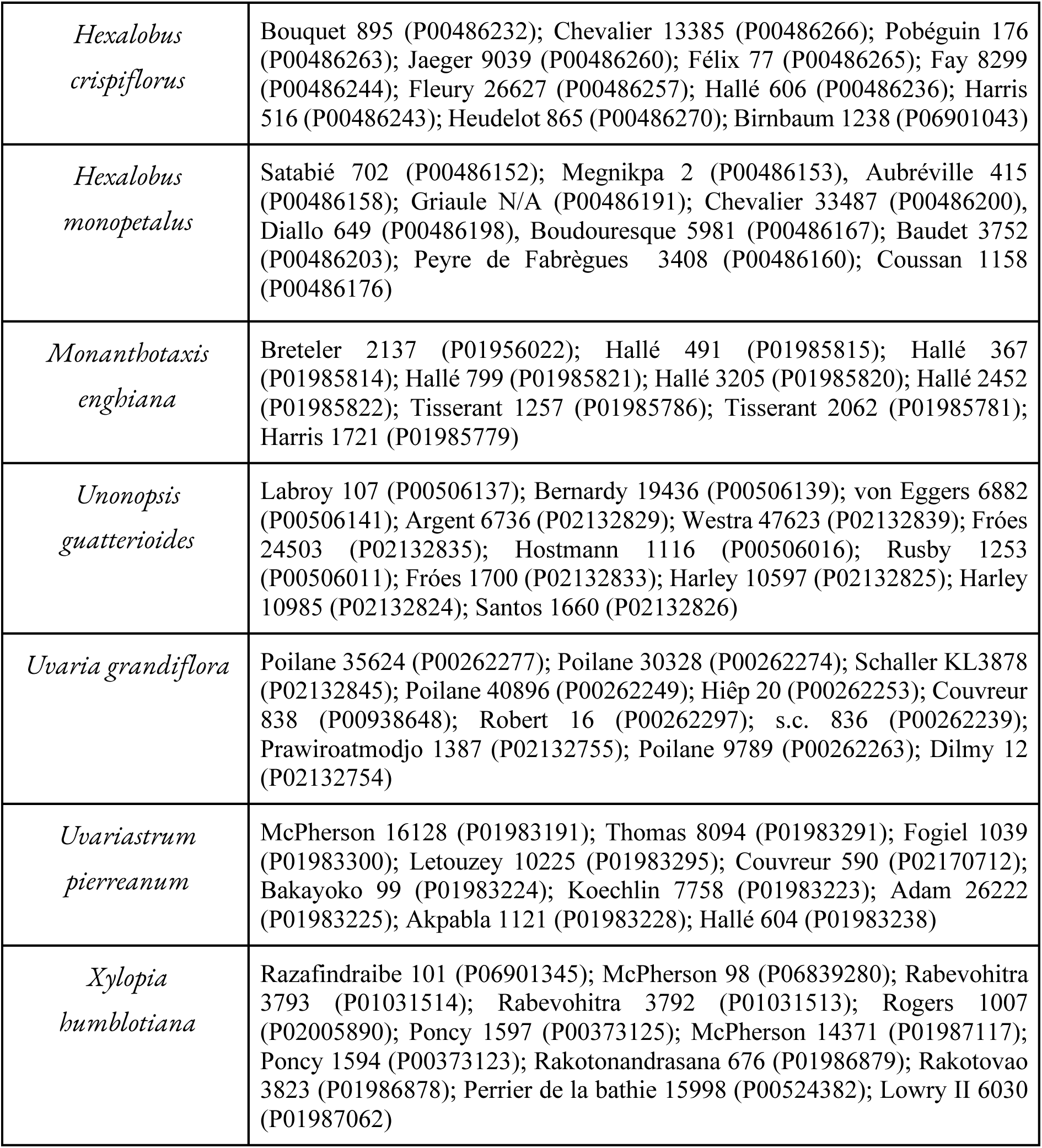
List of herbarium specimens scanned from ID_Paris (barcode in brackets)

**Supp. Figure 1.**
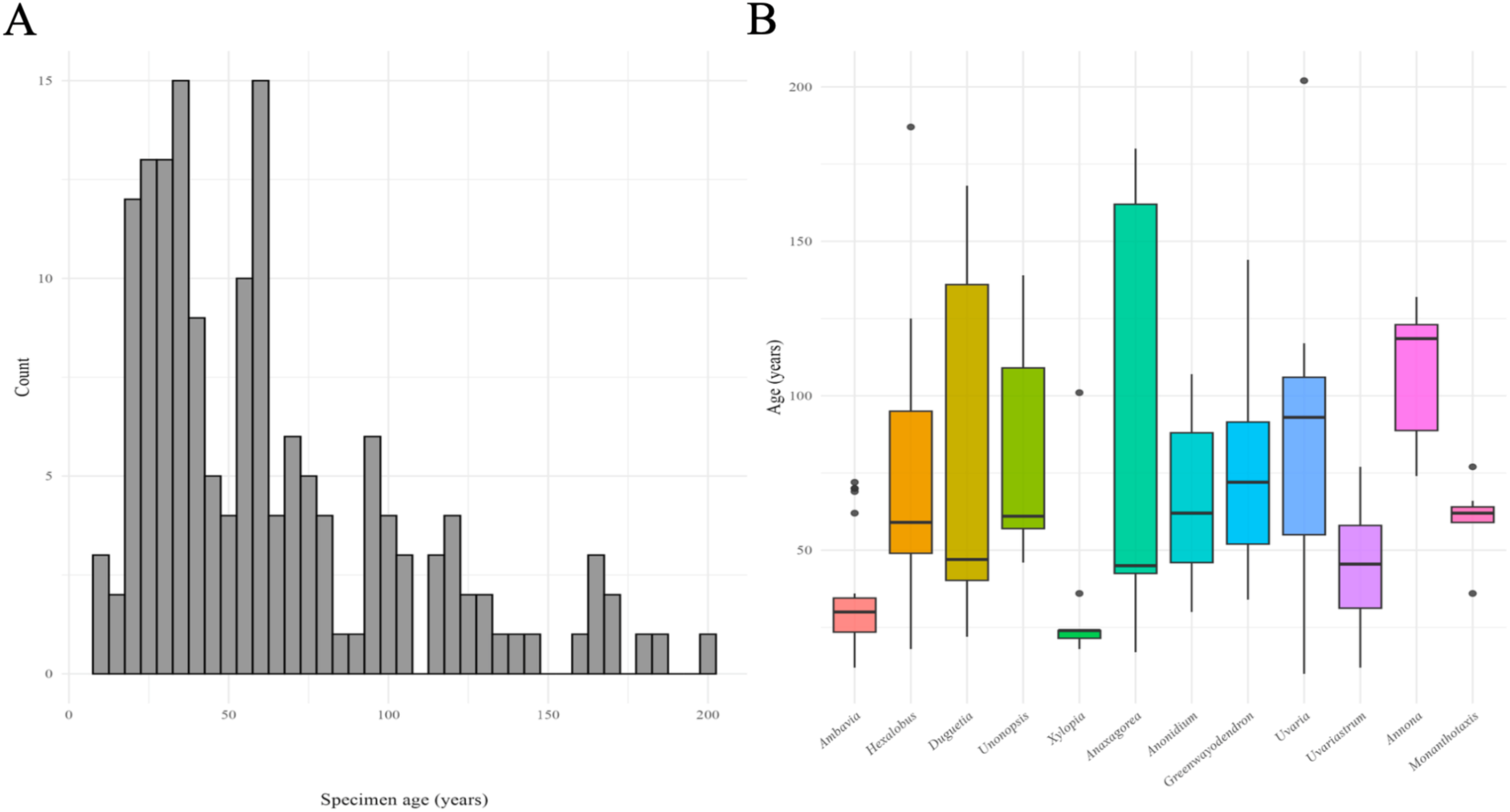
Age distribution of scanned Annonaceae specimens from the Paris herbarium (ID_Paris dataset). **(A)** Histogram of specimen ages (calculated as 2025 minus collection year) showing the distribution of 160 scanned Annonaceae specimens from the Paris herbarium. **(B)** Boxplots of specimen ages grouped by genus. ID_PARIS.

**Supp. Figure 2.**
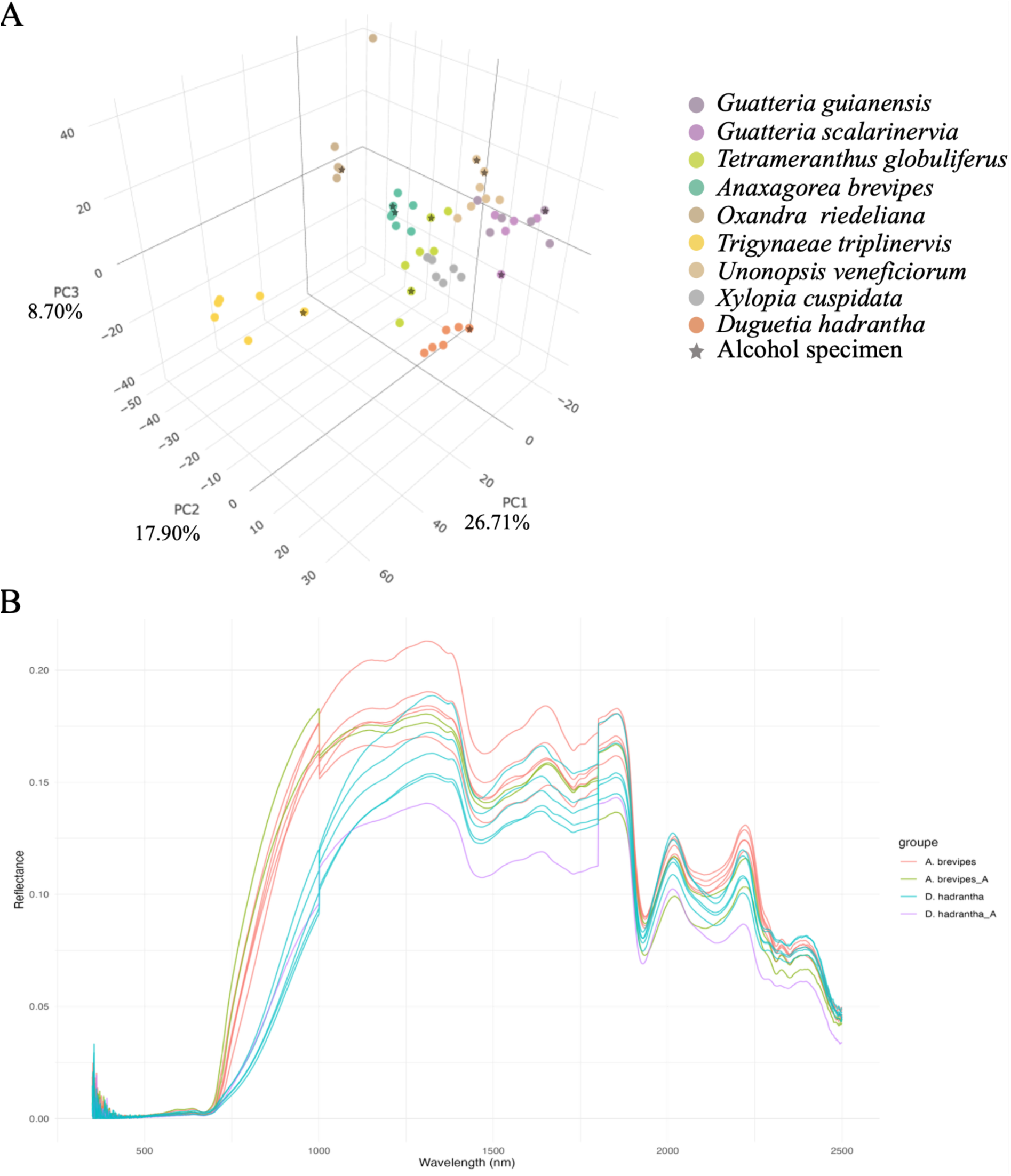
Alcohol preservation effects on leaf spectral data. **(A)** three-dimensional PCA projection of the first three principal components; Find in the Forge repositories a link to the 3D space plot named FS2_3D_plot.Rdata . **(B)** Mean raw reflectance spectra per specimen grouped by species and pre-drying treatment (*Anaxagorea brevipes* and *Duguetia hadrantha* alcohol (_A) vs. directly dried). An averaged spectrum is used to represent the specimen in both analyses. ID_YASUNÍ.

**Supp. Figure 3.**
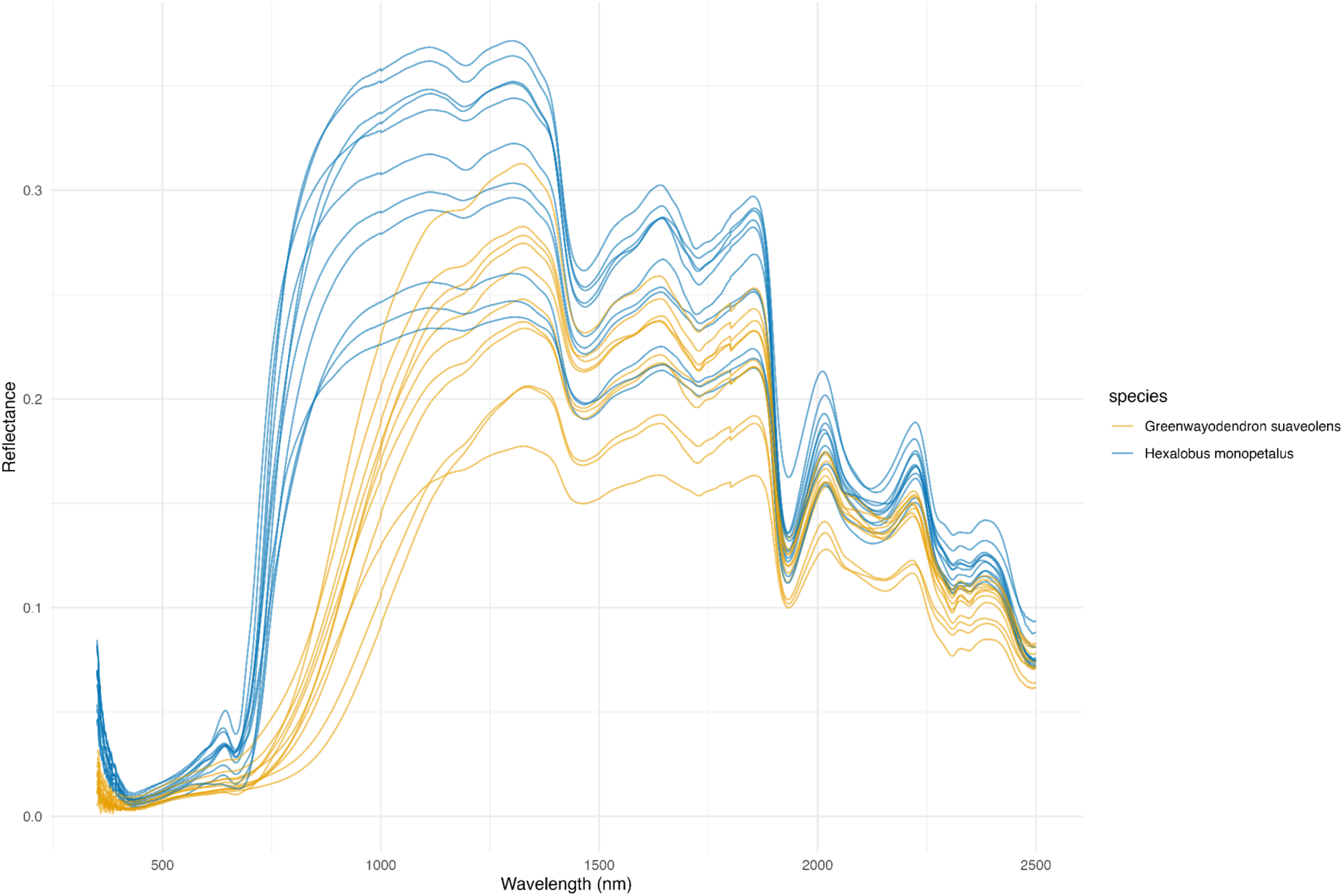
Reflectance spectra from ID_PARIS. Mean raw reflectance spectra per specimen for *Greenwayodendron suaveolens* and *Hexalobus monopetalus*. ID_PARIS.

## Notes

### Competing Interest Statement

The authors have declared no competing interest.

### Summary of Updates

Mainly revisions to the Discussion and improvements in methodological clarity, especially to provide more perspective on the alcohol-related results. Minor revisions were made to some figures for clarity, and one additional supplementary figure was added.

